# Anterior thalamic excitation and feed-forward inhibition of presubicular neurons projecting to medial entorhinal cortex

**DOI:** 10.1101/243022

**Authors:** Mérie Nassar, Jean Simonnet, Li-Wen Huang, Bertrand Mathon, Ivan Cohen, Michael H.K. Bendels, Mathieu Beraneck, Richard Miles, Desdemona Fricker

## Abstract

The presubiculum contains head direction cells that are crucial for spatial navigation. Here, we examined the connectivity and strengths of thalamic inputs to presubicular layer 3 neurons projecting to the medial entorhinal cortex in the mouse. We recorded pairs of projection neurons and interneurons while optogenetically stimulating afferent fibers from the anterior thalamic nuclei (ATN). Thalamic input differentially affects presubicular neurons: layer 3 pyramidal neurons and fast-spiking parvalbumin expressing (PV) interneurons are directly and monosynaptically activated, with depressing dynamics, while somatostatin (SST) expressing interneurons are indirectly excited, during repetitive ATN activity. This arrangement ensures that the thalamic excitation of layer 3 cells is often followed by disynaptic inhibition. Feed-forward inhibition is largely mediated by PV interneurons which have a high probability of connection to presubicular pyramidal cells. Our data point to a specific role of presubicular microcircuits in shaping thalamic head-direction signals transmitted to medial entorhinal cortex: Short-latency PV cell activation may enforce temporally precise head direction tuning during fast turns. However, depression at ATN-PV synapses during repeated activation tends to facilitate pyramidal cell firing when head direction is maintained. Operations performed in presubicular layer 3 circuits seem well-adapted for spatial fine-tuning of head direction signals sent to the medial entorhinal cortex.

**SIGNIFICANCE STATEMENT:** How microcircuits participate in shaping neural inputs is crucial to understanding information processing in the brain. Here, we show how the presubiculum may process thalamic head directional information before transmitting it to the medial entorhinal cortex. Synaptic inputs from the anterior thalamic nuclei (ATN) excite layer 3 pyramidal cells (PC) and parvalbumin (PV) interneurons, which mediate disynaptic feed-forward inhibition. Somatostatin (SST) interneurons are excited indirectly. Presubicular circuits may switch between two regimes according to the angular velocity of head movements. During immobility, SST-PC interactions support maintained head directional firing with attractor-like dynamics. During rapid head turns, in contrast, PV mediated feed-forward inhibition acts to tune the head direction signal transmitted to medial entorhinal cortex.

## INTRODUCTION

The presubiculum is part of the parahippocampal cortex, located between the hippocampus and the entorhinal cortex (Amaral and Witter, 1989; van Strien et al., 2009). It plays a fundamental role in the perception of spatial orientation. Most presubicular neurons of layer 3 and some in deeper layers signal head direction (Boccara et al., 2010; Preston-Ferrer et al., 2016). These cells fire persistently when the head of an animal is oriented in a specific direction. Head direction information originates subcortically, mostly from vestibular inputs. The anterior thalamic nucleus (ATN) relays these signals to the presubiculum (Taube, 2007; van Groen and Wyss, 1990; Peyrache et al., 2015). Lesions of the anterior thalamus abolish head direction firing and impair grid cell signals in parahippocampal cortex (Goodridge and Taube, 1997; Winter et al., 2015). In the presubiculum thalamic head direction signals (Taube et al., 1995) are integrated with visual information from visual (Vogt and Miller, 1983) and retrosplenial cortices (van Groen and Wyss, 1990). Neurons of the ATN also contribute to spatial firing of grid cells of medial entorhinal cortex (MEC) (McNaughton et al., 2006; Langston et al., 2010, Winter et al., 2015). The presubicular projection may therefore transmit ATN derived head direction signals to the medial entorhinal grid cell system (Rowland et al., 2013; Preston-Ferrer et al., 2016). However it is unclear whether thalamic axons contact presubicular cells that project to the MEC.

Recent work has revealed some properties of circuits involving presubicular principal cells and interneurons (Nassar et al., 2015; Simonnet et al., 2013; Peng et al., 2017). Local interactions between pyramidal cells and Somatostatin (SST) expressing Martinotti cells may help sustain firing in head direction cells (Simonnet et al., 2017). The role of fast spiking parvalbumin positive (PV) interneurons in processing thalamic head directional information remains to be clarified, however. Here we examine the hypothesis that these interneurons mediate a feed-forward inhibitory control of head direction signals transmitted to the grid cell system. We used electrophysiology, optogenetics and retrograde labeling to ask how examine presubicular PV cells and pyramidal cells process excitatory inputs from the ATN. Our data shows ATN axons directly contact pyramidal MEC projecting neurons and PV interneurons of layer 3. PV interneurons mediate feed-forward inhibition, and their recruitment is strongest at the onset of ATN spiking.

## MATERIALS AND METHODS

### Animals

Experiments were performed on male and female offspring of male Pvalb-Cre mice (Jax 008069; Hippenmeyer et al., 2005) or Sst-IRES-Cre mice (Jax 013044; Taniguchi et al., 2011) crossed with the Ai14 Cre reporter line (Jax 007914; Madisen et al., 2010). Cre-mediated recombination resulted in the expression of red fluorescent tdTomato labeling in Pvalb-Cre or SST-Cre neurons. Wild-type Bl6 and Pvalb-Cre males were used for double injections. Animal care and use conformed to the European Community Council Directive of 22 September 2010 (2010/63/EU) and French law (87/848). Our study was approved by the ethics committee Charles Darwin N○5 and the French Ministry for Research (n° 01025.02).

### Viral vectors

Viral vectors were injected to induce expression of channelrhodopsin and halorhodopsin. AAV2.hSyn.hChR2(H134R)-eYFP.WPRE.hGH (Penn Vector Core, Addgene 26973P) with serotypes 5 or 9, contained an enhanced ChR2-EYFP fusion gene, driven by a hSynapsin promoter. We also used AAV1.EF1a.DIO.eNpHR3.0-eYFP.WPRE.Hgh (Penn Vector Core, Addgene 26966) an adeno-associated virus serotype 1 carrying Cre-inducible halorhodopsin-3.0 (eNpHR3.0)-enhanced yellow fluorescent protein (eYFP) transgene driven by EF1a promoter for optogenetic inhibition. Viral vectors were stored in aliquots at −80°C until use.

### Stereotaxic surgery

Mice, of age 25-35 days, were anesthetized intraperitoneally with ketamine hydrochloride and xylazine (100 and 15 mg.kg^−1^, respectively) and placed in a stereotaxic frame. Unilateral viral injections were performed as previously (Mathon et al., 2015). AAV injections were targeted to Anterior-thalamic nuclei (ATN) using coordinates from bregma of lateral, +0.75 mm; posterior, −0.82 mm; and depth, 3.2 mm. Injections were made with a Hamilton syringe using a syringe Pump Controller (Harvard Apparatus, Pump 11 elite) at 60 nL.min^−1^ to deliver volumes of 150-300 nL hSyn-ChR2 AAV2/5 or AAV2/9 vectors. Leakage from an injection site to surrounding tissue was minimized by withdrawing the pipette slowly after a delay of 5 min. Maximal transgene expression occurred 2 weeks after AVV2/9 and 3 weeks after AAV2/5 injection. Viral titers for AAV2/5 and AAV2/9 vectors were respectively 1.3^e^10^13^ and 3.39^e^10^13^ virus particles.ml^−1^. Since light-evoked responses were similar for ChR2 expression by either AAV serotype, results were pooled. We also made double injections of AAV in ATN and a retrograde tracer in MEC. Presubicular projecting neurons were labeled retrogradely by red fluorescent latex microspheres (Lumafluor; volume 300 nL) injected in the ipsilateral MEC at coordinates from bregma for MEC of lateral, +3 mm; posterior, −4.7 mm; and depth, 4 mm. Finally, we made double injections of AAV-ChR2 in ATN and AAV-eNpHR3.0 in the presubiculum of Pvalb-Cre mice to express halorhodpsin selectively in PV interneurons. AAV-EF1a-DI0-eNpHR3.0-EYFP was injected (final volume 350 nL) in the presubiculum at coordinates from bregma of lateral +2 mm; posterior −4.06 mm; and depth 2.15 mm. The viral titer was 1.09^e^13 particles.ml^−1^ and 5 weeks was needed for full expression.

### Preparation of brain slices

Slices containing the hippocampus, subicular complex and entorhinal cortex were prepared 2–5 weeks after virus and/or tracer injection. Animals were anesthetized with ketamine hydrochloride and xylazine (100 and 15 mg.kg^−1^, respectively). They were perfused through the heart with a solution of 125 NaCl, 25 sucrose, 2.5 KCl, 25 NaHCO_3_, 1.25 NaH_2_PO_4_, 2.5 D-glucose, 0.1 CaCl_2_, 7 MgCl_2_ (in mM) cooled to 4°C and equilibrated with 5% CO_2_ in O_2_. The brain was removed, and horizontal, 300-320 μm thick sections cut in the same solution with a vibratome (Leica VT1000S). They were stored for at least 1 h at 22–25°C in a chamber filled with ACSF containing (in mM): 124 NaCl, 2.5 KCl, 26 NaHCO_3_, 1 NaH_2_PO_4_, 2 CaCl_2_, 2 MgCl_2_, and 11 D-glucose, bubbled with 5% CO_2_ in O_2_ (pH 7.3, 305–315 mOsm/L). Slices were then transferred to a recording chamber (volume 2–3 ml, temperature 33–35°C) mounted on a BX51WI microscope (Olympus, France). Experiments were discontinued if transfection at the injection site was weak or if leakage had occurred.

### Whole-Cell Patch-Clamp Recordings

Recordings were made with glass pipettes pulled using a Brown-Flaming electrode puller (Sutter Instruments) from borosilicate glass of external diameter 1.5 mm (Clark Capillary Glass, Harvard Apparatus). Electrodes were filled with a solution containing (in mM): 135 K-gluconate, 1.2 KCl, 10 HEPES, 0.2 ethylene glycol tetra-acetic acid (EGTA), 2 MgCl_2_, 4 MgATP, 0.4 Tris-GTP, 10 Na_2_-phosphocreatine and 2.7–7.1 biocytin. QX314 bromide (2 mM; Tocris) a blocker of voltage-activated Na^+^ channels was added to the pipette solution when inhibitory (I) and excitatory (E) synaptic currents were measured at depolarized holding potentials (Fig. 3). The pH of the pipette solution was adjusted to 7.3 with KOH and the osmolarity was 290 mOsm. Electrode resistance was 4–8 MOhm. Fluorescent PV or SST cells were identified using LED illumination with appropriate emission/excitation filters (OptoLED, Cairn Research, Faversham, UK) using a Luca CCD Camera (Andor). Whole-cell current-clamp recordings were made using a MultiClamp 700B amplifier and pCLAMP software (Molecular Devices, Union City, CA, USA). Membrane potential signals were filtered at 6 kHz and digitized at 20–50 kHz. An estimated junction potential of 15 mV was not corrected. Pyramidal cells were identified as non-fluorescent regular-spiking neurons, Parvalbumin- or Somatostatin-expressing interneurons were identified as red fluorescent cells in tissue from PvalbCre::tdTomato and Sst-Cre::tdTomato mice respectively. Neuronal electrical properties were measured from hyperpolarizing and depolarizing step current injections and analyzed with Axograph X and routines written in MATLAB, as previously (Nassar et al., 2015; Table 1). PV interneurons recorded from PvalbCre::tdTomator mice consistently fired action potentials with fast kinetics. We have suggested that labelled SST interneurons of Sst-Cre::tdTomato mice are heterogenous (Nassar et al., 2015). This work was limited to labelled SST cells with a low threshold firing pattern (LTS).

**Table 1.**
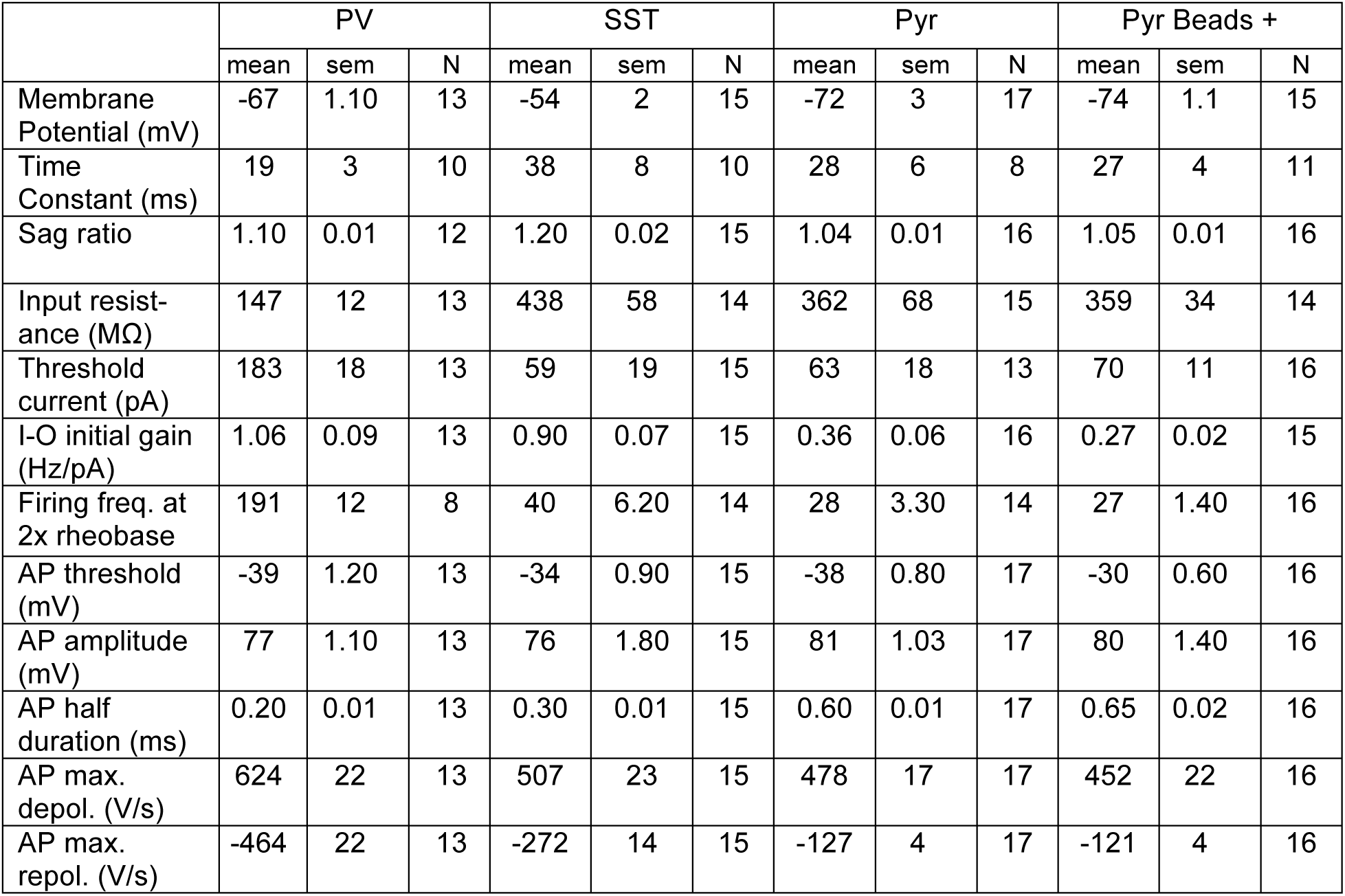
Intrinsic properties of layer 3 presubicular neurons.

|  | PV |  |  | SST |  |  | Pyr |  |  | Pyr Beads + |  |  |
| --- | --- | --- | --- | --- | --- | --- | --- | --- | --- | --- | --- | --- |
|  | mean | sem | N | mean | sem | N | mean | sem | N | mean | sem | N |
| Membrane Potential (mV) | -67 | 1.10 | 13 | -54 | 2 | 15 | -72 | 3 | 17 | -74 | 1.1 | 15 |
| Time Constant (ms) | 19 | 3 | 10 | 38 | 8 | 10 | 28 | 6 | 8 | 27 | 4 | 11 |
| Sag ratio | 1.10 | 0.01 | 12 | 1.20 | 0.02 | 15 | 1.04 | 0.01 | 16 | 1.05 | 0.01 | 16 |
| Input resistance (MΩ) | 147 | 12 | 13 | 438 | 58 | 14 | 362 | 68 | 15 | 359 | 34 | 14 |
| Threshold current (pA) | 183 | 18 | 13 | 59 | 19 | 15 | 63 | 18 | 13 | 70 | 11 | 16 |
| I-O initial gain (Hz/pA) | 1.06 | 0.09 | 13 | 0.90 | 0.07 | 15 | 0.36 | 0.06 | 16 | 0.27 | 0.02 | 15 |
| Firing freq. at 2x rheobase | 191 | 12 | 8 | 40 | 6.20 | 14 | 28 | 3.30 | 14 | 27 | 1.40 | 16 |
| AP threshold (mV) | -39 | 1.20 | 13 | -34 | 0.90 | 15 | -38 | 0.80 | 17 | -30 | 0.60 | 16 |
| AP amplitude (mV) | 77 | 1.10 | 13 | 76 | 1.80 | 15 | 81 | 1.03 | 17 | 80 | 1.40 | 16 |
| AP half duration (ms) | 0.20 | 0.01 | 13 | 0.30 | 0.01 | 15 | 0.60 | 0.01 | 17 | 0.65 | 0.02 | 16 |
| AP max. depol. (V/s) | 624 | 22 | 13 | 507 | 23 | 15 | 478 | 17 | 17 | 452 | 22 | 16 |
| AP max. repol. (V/s) | -464 | 22 | 13 | -272 | 14 | 15 | -127 | 4 | 17 | -121 | 4 | 16 |

### Optical stimulation

ChR2-expressing ATN axons, were stimulated optically with either a single-wavelength LED system (470 nm; Cairn OptoLED) or laser illumination (405 nm; LuxX, Omicron), connected to the microscope via a custom made triple port (Cairn). Synaptic responses of presubicular neurons, usually simultaneously recorded pyramidal cells and interneurons, were recorded in whole-cell current- or voltage-clamp modes. LED illumination gave a 200 μm-diameter spot with a 60X 1.0 NA plan-Apochromat objective. Flashes of 0.5–20 ms were delivered close to a recorded somata. We typically used low intensity stimulation of 0.1-0.5 mW, set to induce near threshold responses, reducing network activation after ATN axon stimulation. Excitatory and inhibitory post-synaptic responses were evoked by 10 Hz and 30 Hz stimulus trains (light pulse duration, 0.5 ms). We tested for monosynaptic excitation by stimulating in the presence of TTX (1 μM; Tocris) to block polysynaptic excitation, and 100 μM 4-AP (Sigma) to enhance axonal depolarization (Mao et al., 2011; Cruikshank et al., 2010). NBQX (10 μm, Tocris), D-AP5 (100 μm, Tocris), Gabazine (10 μm, Tocris) and CGP 52432 (10 μM, Tocris) were used to block AMPA, NMDA, GABA_A_ and GABA_B_ receptors respectively.

Inhibition of PV interneurons during photostimulation of the thalamic pathway, was achieved by combining a yellow LED (572 nm) photo-activation of eNpHR3.0 in PV neurons, with a blue light (470 nm) illumination in the lightpath. Light intensity at the sample was set at 0.3 mW. The yellow light pulse, of 20 ms duration, was triggered 5 ms before a blue light pulse of 0.5 ms duration so that PV interneurons were inhibited before and during thalamic axon stimulation. For each injected mouse, the efficacy of yellow LED-triggered hyperpolarization was verified in at least one PV interneuron.

The excitatory and inhibitory effects of stimulating thalamic fibers at different presubicular sites were explored in layer 3 pyramidal cells clamped to potentials between −30 mV and 0 mV. In this way biphasic synaptic events could be detected as a focused 405 nm laser beam (LuxX, Omicron) was scanned over different presubicular sites (cf. Fig. 3F). After acquiring an initial image with a 4X objective, we switched to a 60X objective for focal point illumination (spot size 5-10 μm). A motorized platform (Luigs and Neumann, Ratingen, Germany) moved the stage across a scanning field as we explored a hexagonal grid of optical stimulation sites (custom-software, Morgentau Solutions, Munich, Germany; Bendels et al., 2008; Beed et al., 2010). 40 to 80 stimulation sites at 40 μm separation were tested.

The spatial resolution of laser illumination is defined by the single photon illumination cone and diffraction limited by light scattering in the slice (Trigo et al., 2009). A light spot of ~ 10 μm diameter in 100 μM pyranine (HPTS) solution was measured at the focus using the Luca CCD camera. The light was set to deliver 0.5-2 ms pulses at each stimulation site. Laser power was adjusted to 0.4-2 mW, to record EPSC-IPSC sequences resulting from activation of few fibers. Grids were repeated three times for each recorded neuron.

We attempted to limit variability from differences in viral vector uptake/expression. First, records were made exclusively from slices where ChR2-eYFP expression was visible at 4X magnification. Second, recordings were only made from superficial layer 3 where ChR2 expression was high. Third, we only used data from animals where significant excitatory responses were initiated in pyramidal cells in control conditions and after TTX/4AP application. Finally, double-recordings from neighboring pyramidal cells and PV or SST interneurons let us normalize responses evoked in an interneuron to those induced by the same photo-stimulation in a nearby pyramidal cell.

### Data analysis

Peak amplitudes of averaged light-evoked PSCs (n = 30) were measured for PV and pyramidal cells. Failures and additional or later peaks were detected for EPSCs evoked in SST interneurons but not PV interneurons or pyramidal cells. We also measured the amplitude at the first peak of individual traces. Synaptic event amplitudes of > 3 times the standard deviation of the noise were considered to result from direct and/or disynaptic input(s). Success rate was determined as the number of detected EPSCs, divided by the total number of stimuli. EPSCs were recorded at holding potentials close to −60 mV and IPSCs near 0 mV. EPSC latency was calculated from the stimulus to 5 % of the rising amplitude of the evoked current from individual traces and then averaged. Latency was measured to the first peak when multiple EPSC peaks were evoked. Short-term synaptic dynamics were analysed by normalizing responses to each stimulus to the amplitude of the first event. IPSCs were recorded at 0 mV, in the presence of D-AP5, and were abolished by gabazine. IPSC onset was measured at 5% of the rising amplitude of the peak outward current. Onset jitter was defined as the SD of latencies to this 5% amplitude point from repeated single sweeps. Paired-pulse ratio (PPR) was defined as the ratio of the amplitude of the second to the first postsynaptic response to dual stimuli. It was calculated from averages of responses to ten stimuli. Post-synaptic responses were recorded at resting membrane potential in current-clamp, except for SST interneurons which were held at membrane potentials near −50-55 mV to suppress firing. Action potential latencies were calculated as the delay from stimulus onset to the peak of the action potential. Spiking probability corresponded to the number of action potentials generated during a train of 10 stimulations for a given intensity, divided by 10. Input resistance was monitored by small current injection steps (-50 pA). Recordings were discarded if changes exceeded 20%.

### Paired recordings

Whole-cell recordings of pairs of synaptically connected neurons were made in voltage clamp mode. Depolarizing voltage injections (1–2 ms, 100-200 mV) were used to elicit single action currents in a presynaptic neuron. Unitary uIPSCs were recorded at −50 mV and uEPSCs at −70 mV holding potential in the postsynaptic cell. Excitatory and inhibitory postsynaptic currents were automatically detected and measured from filtered records (low-pass 1-1.5 KHz) with a threshold of 4-6 pA for EPSCs and IPSCs. The properties of unitary IPSCs and EPSCs were determined from 30–60 traces. Peak amplitude was defined as the mean of all responses including failures (amplitude close to 0 pA). Synaptic latency was estimated as the time between the peak of the presynaptic action current and 5% of the peak amplitude of a unitary uIPSC. Rise time of synaptic currents was defined as the time from 20 to 80% of their peak amplitude. Synaptic event decay times were measured between peak and 50% of the peak. For analyses of short-term plasticity, PSCs in a train were normalized to the amplitude of the first EPSC or IPSC. Synaptic transfer rate was calculated as the number of detected post-synaptic events divided by the number of presynaptic spikes. Failure rate was 1 - transfer rate.

### Image acquisition and Analysis

Stained slices were visualized with a QImaging Retiga EXI camera (Qimaging Surrey, BC, Canada), and scanned with an Optigrid II (Thales Optem, Qioptik, Rochester, NY, USA) on an inverted Olympus IX81 microscope to acquire structured images. Stacks of 50–80 images (z-step, 0.7 μm) were acquired with an oil immersion objective (20x, NA 0.9). Presubicular layers and borders were defined from cytoarchitectonic features defined from nuclear (DAPI) staining. Higher resolution images (Fig. 2B) were obtained using an Olympus FV-1000 confocal microscope to acquire z-stacks at 0.3 μm. Blue, green, red, fluorescent signals were acquired sequentially and images adjusted for contrast and brightness with Image J (NIH).

### Morphological 3D reconstructions

Neurons were filled with 0.3% biocytin (3 mg/mL) to reveal the morphology of recorded cells (Nassar et al., 2015; Simonnet et al., 2013). Neuronal form was derived from z-stacks of acquired images (Neurolucida; Microbrightfield, Williston, VT, USA).

### Statistics

Signals were analyzed with AxoGraphX, and custom written software (Labview, National Instruments; MATLAB, The Mathwork). Algorithms to detect action potentials and measure active and passive neuronal properties are as described (Simonnet et al., 2013; Nassar et al., 2015). Results are given as mean ± SEM (n=number of cells, slices, or animals as indicated). Statistical analysis was performed with Prism (GraphPad Software, Inc.), MATLAB (The Mathwork) and R. The Wilcoxon signed rank test for matched pairs was used to compare non-parametric data in matched samples. A Kruskal–Wallis one-way analysis of variance (ANOVA) test was followed by Dunn’s *post hoc* test to compare more than two groups. The Anderson-Darling two-sample procedure (alpha=0.05) was used to compare non-parametric data with different distributions. The Benjamini and Hochberg procedure was used to correct p-values for multiple testing. Significance levels are indicated as *p* values.

## RESULTS

### Selective ChR2 expression in the anterior thalamic nuclei

In order to study the functional connectivity of the anterior thalamic nuclei (ATN) to the presubiculum, we made unilateral injections of adeno-associated virus (AAV) to express light-gated Channelrhodopsin-2 fused to green fluorescent protein (ChR2-eYFP) in ATN neurons. After two to three weeks of incubation, horizontal slices were prepared, and the thalamic injection site was examined. Fig. 1A, B shows an example of ChR2-eYFP expression in the ATN. Labeled (eYFP) thalamic axons innervated layers 1 and 3 of the presubiculum more densely than layer 2. Thalamic afferents did not project to deep presubicular layers, adjacent subiculum or entorhinal cortex (Fig. 1C, D). Recording from ChR2-eYFP expressing thalamic neurons in slices revealed typical rebound bursting (Fig. 1E). Illumination with blue light in the presence of the glutamatergic antagonists CNQX and D-AP5 depolarized these cells and induced firing in current clamp records (Fig. 1F, upper). In voltage clamp mode, an inward ChR2-mediated photocurrent was detected (Fig. 1F, lower). ChR2-eYFP expressing thalamic neurons responded to light with latencies less than 0.1 ms (n = 4 neurons from 2 mice).

**Figure 1:**
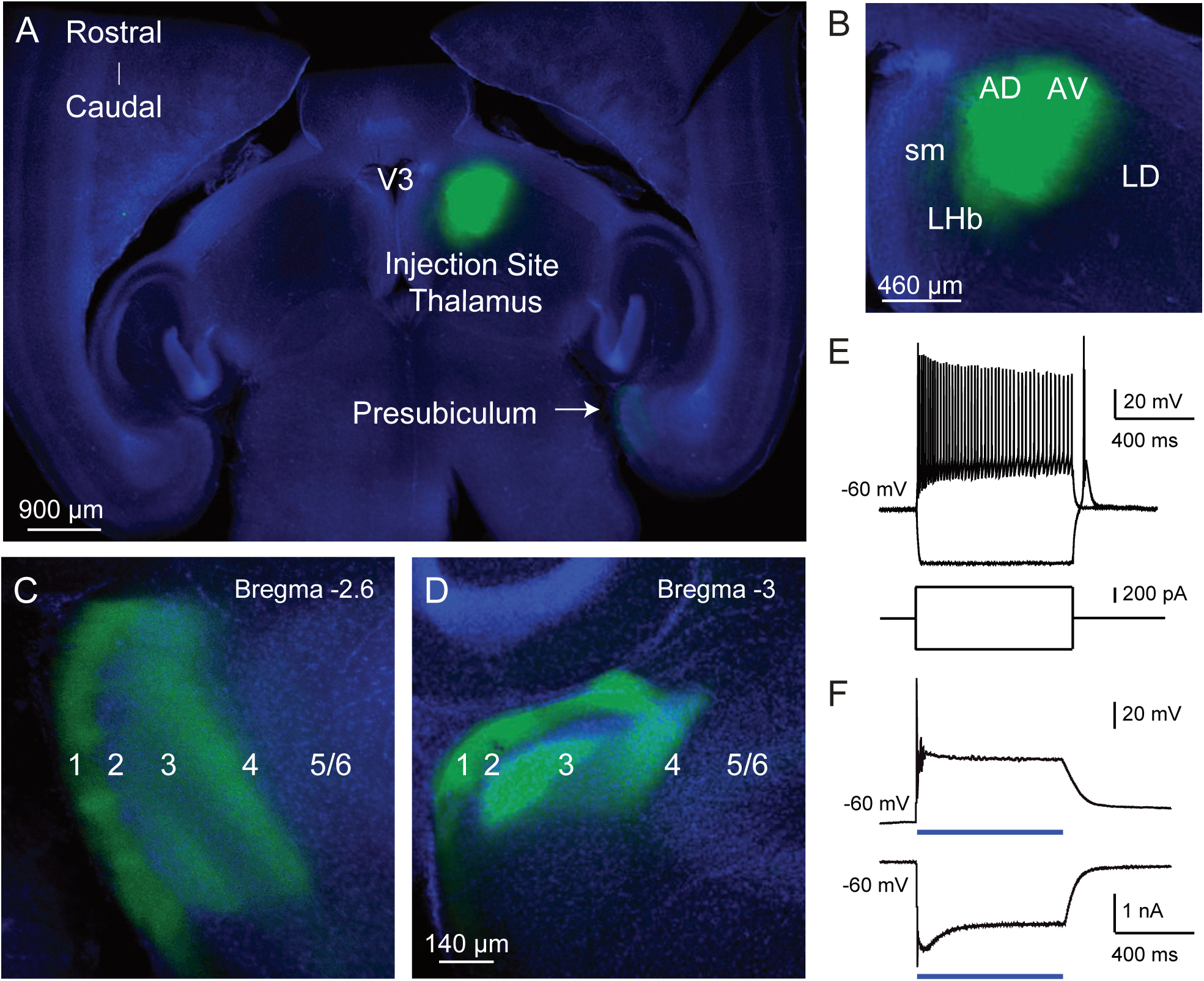
ChR2-eYFP expression in the anterior thalamic nuclei and in their axon terminals in the presubiculum. (A) Mouse brain injected with AAV5-ChR2-eYFP (green) in the anterior thalamic nuclei (ATN). 320 μm-thick horizontal section with DAPI staining in blue. (B) ChR2-eYFP expression at the injection site. AD, antero-dorsal nucleus; AV, antero-ventral nucleus; LD, latero-dorsal nucleus; sm, stria medullaris; V3, third ventricle; LHb, lateral habenula. (C), (D) ChR2-eYFP expressing axon terminals in the presubiculum at different dorso-ventral levels, Bregma −2.6 and −3 mm. Presubicular layers 1 to 6 are indicated. (E) Firing of a thalamic cell induced by a two-fold rheobase depolarizing current. Hyperpolarization induced by a negative current pulse is followed by a rebound burst. (F) Top, blue light stimulation (0.1 mW, 300 ms) evoked spikes and depolarization block in the thalamic cell. Bottom, a ChR2-mediated photocurrent recorded in voltage-clamp in response to the same photostimulation. The blue bar indicates the light pulse. Recordings made in the presence of CNQX and D-AP5.

### ATN axon terminals directly contact MEC projecting pyramidal neurons

We asked whether ATN axons innervate MEC projecting pyramidal cells (PCs), by combining retrograde labeling of MEC-projecting cells in the presubiculum with the viral expression of ChR2-eYFP in thalamic axons (Fig. 2A,B). Red retrobeads were observed in neuronal somata located in the presubicular superficial layer and a few deeper neurons (Fig. 2B, C; cf. Huang et al., 2017), indicating that these project to the MEC. In *ex vivo* slice recordings, light stimulation of ATN axons induced excitatory postsynaptic currents (EPSCs) in all retrobead-labeled layer 3 neurons with somata surrounded by channelrhodopsin-expressing ATN axons (Fig. 2D,F, same neuron as in 2C; low intensity LED stimulation, 0.1 − 0.5 mW). The mean delay to EPSC onset was 2 ± 0.15 ms (n = 11, Fig. 2E), compatible with monosynaptic activation. Evoked EPSC amplitudes varied for different neurons in different slices (289 ± 43 pA, mean ± sem, n = 11 neurons from 5 animals; Fig. 2E). In current-clamp recordings, ATN fiber stimulation initiated action potentials in bead-labeled MEC-projecting neurons at resting membrane potential (Fig. 2F, stimulations repeated at 30 Hz). Action potentials were abolished by TTX (1 μm) and 4-AP (100 μM), but light stimuli continued to induce excitatory postsynaptic potentials (EPSPs), indicating that post-synaptic cells were contacted directly by synapses of ChR2-expressing ATN axons (n = 11; cf. Mao et al., 2011). EPSCs were suppressed by bath application of AMPA and NMDA receptor antagonists (Fig. 2G, n = 3 neurons from three retrobead labeled mice). The glutamatergic nature of thalamic excitation was confirmed in four additional layer 3 pyramidal neurons from three mice that had not received retrobead injections.

**Figure 2:**
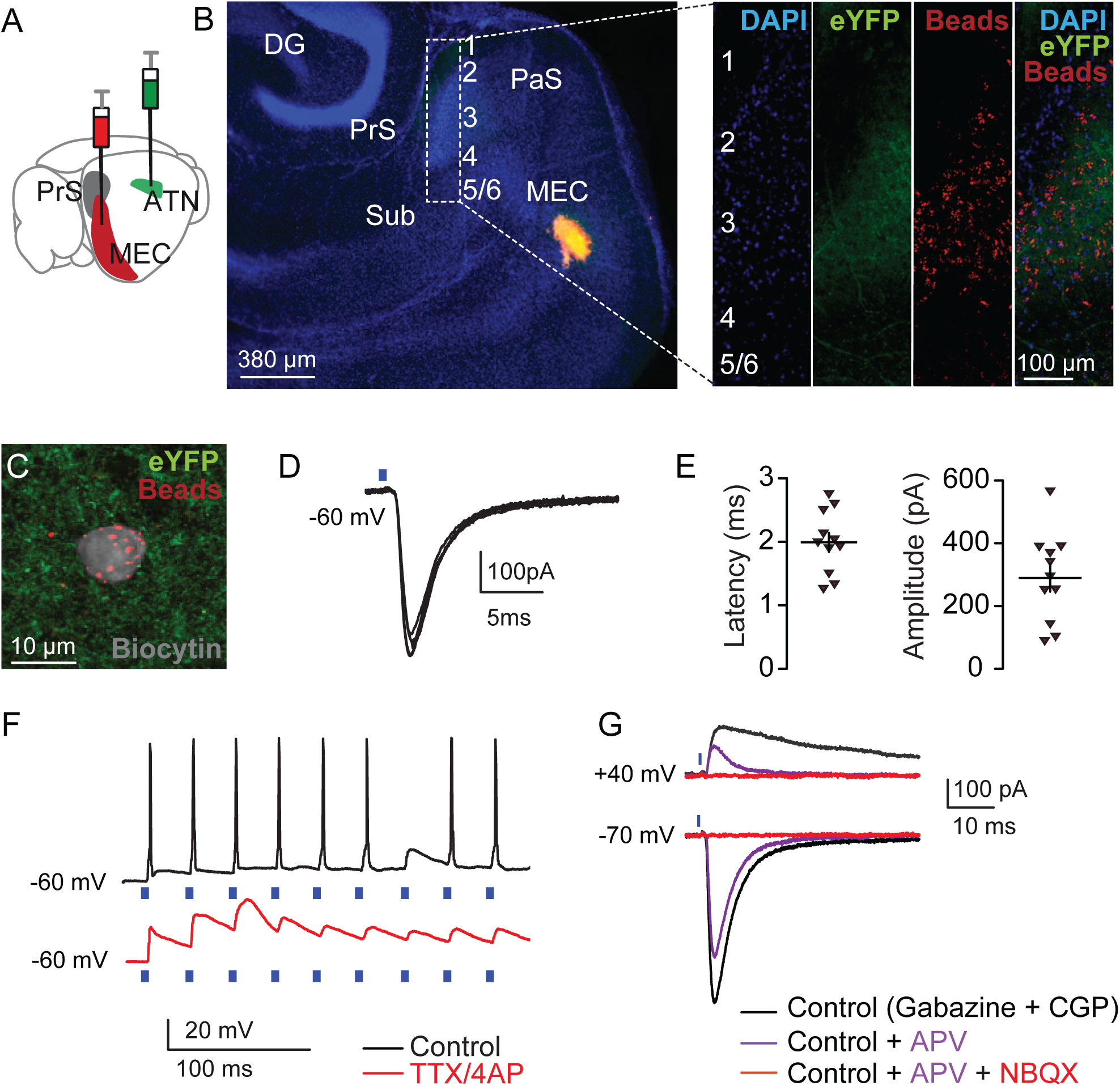
ATN axons contact MEC projecting neurons. (A) Schematic showing unilateral injection sites for AAV5-ChR2 in ATN (green) and retrobeads in the MEC (red). (B) Left, horizontal section of the parahippocampal formation with retrobeads injected in the MEC in orange. DAPI staining in blue. Thalamic axons expressing ChR2-eYFP target the presubiculum. DG: dentate gyrus, PrS: presubiculum, Sub: subiculum, PaS: parasubiculum, MEC: medial entorhinal cortex. Right, magnification of the presubiculum (inset in B) stained with DAPI (blue), ChR2-eYFP expressing thalamic axons (green), retrobeads (red) and a merged image (right). (C) Confocal image of the soma of a biocytin-filled neuron in superficial layer 3. The soma (grey) contains red retrobeads and is surrounded by thalamic axons (green). (D) Light-evoked EPSCs in the same neuron. (E) Average EPSC latencies (left) and amplitudes (right) from 11 bead-labeled neurons. Mean ± SEM are indicated by horizontal and vertical lines. (F) Light stimuli repeated at 30 Hz initiated action potential firing (upper trace, black; same neuron as in (C). TTX (1 μM) and 4-AP (100 μM) abolished spikes, while direct EPSPs persisted (bottom trace, red). (G) Light evoked currents at holding potentials of +40 and at -70 mV revealed glutamatergic EPSCs with a NMDA and AMPA receptor mediated component. Recordings in the presence of 10 Gabazine and 10 μm CGP 52432. EPSCs were entirely abolished by 100 μM APV and 10 NBQX.

### Activation of ATN inputs drives an excitation/inhibition sequence in the presubiculum

When presubicular layer 3 pyramidal neurons were voltage-clamped at 0 mV, photostimulation of ChR2-expressing thalamic fibers initiated biphasic synaptic currents. (Fig. 3A, B). An initial excitatory inward current (EPSC) was rapidly followed by an inhibitory outward current (IPSC). Following whole-field LED stimulation, both EPSCs and IPSCs were elicited in all layer 3 pyramidal neurons tested (n = 9 PCs from 4 animals), including retrobead labeled MEC projecting PCs (n = 3 PCs from 3 animals). IPSCs were abolished by both the GABA receptor antagonist Gabazine and the AMPA receptor antagonist NBQX (Fig. 3A, B). IPSC onset latencies were significantly longer (Fig. 3C; 3.3 ± 0.1 ms; mean ± sem; n = 12 from 5 animals; p < 0.0001, Wilcoxon signed rank test) than those of the EPSCs (1.7 ± 0.1 ms; mean ± sem; n = 12 from 5 animals). In PCs, the delay between EPSC and IPSC onset was 1.6 ± 0.1 ms. Onset latency jitter for EPSCs was significantly less than that of IPSCs (EPSCs, 0.12 ± 0.03 ms; IPSCs 0.23 ± 0.03 ms; mean ± sem; n = 12 cells from 5 animals; p < 0.05, Wilcoxon signed-rank test, Fig. 3D). These data show that photostimulation of ATN afferents can evoke a direct glutamatergic excitation followed by a disynaptic inhibition, compatible with thalamic-induced feed-forward inhibition of layer 3 presubicular PCs (Fig. 3E top).

**Figure 3:**
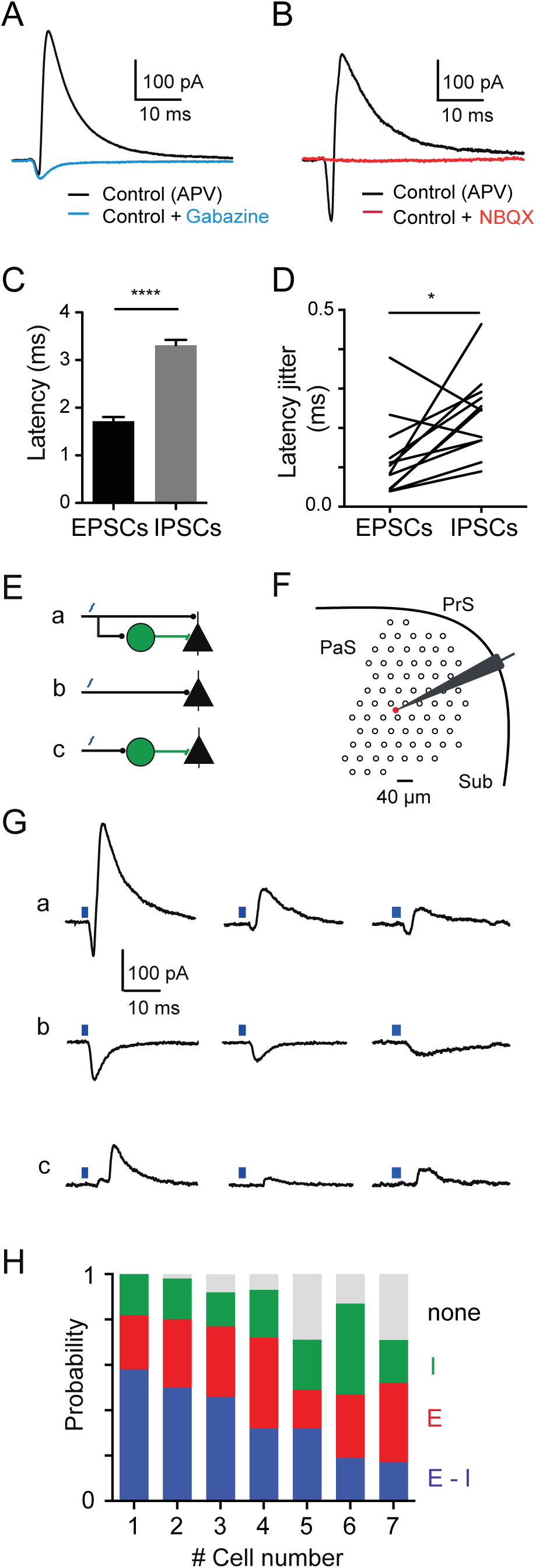
ATN driven sequences of excitation-inhibition vs. unbalanced excitation or inhibition. (A), (B) ATN-evoked currents recorded from presubicular pyramidal neurons at 0 mV holding potential, in the presence of APV. The current was biphasic, with an initial inward current followed by an outward current. (A) The outward current, mediated by GABAA receptors, was entirely blocked by gabazine. (B) NBQX abolished both the inward current and the disynaptic GABAergic component. (C), (D) Disynaptic IPSCs induced by ATN stimuli have longer onset latencies and more jitter than the ATN-driven EPSCs. * p < 0.05, **** p < 0.001, Wilcoxon matched-pairs signed rank test. Values are given as mean ± SEM. (E) different possibilities of afferent connectivity: a pyramidal neuron may receive either (a) excitation and feedforward inhibition, (b) excitation only or (c) inhibition only. (F) Schematic of the experiment showing the presubiculum (PrS) with the recording pipette and the pyramidal cell soma (red), and the grid of laser light stimulation sites (black circles, spacing 40 μm). PaS, parasubiculum; Sub, Subiculum. (G) a, sequences of excitatory-inhibitory responses, b, excitatory responses only, or c, inhibitory responses only. Recordings from a same pyramidal neuron for different laser light stimulation sites across the presubiculum. (H) Probability to elicit excitatory-inhibitory responses (E-I, blue), excitatory responses only (E, red), or disynaptic inhibitory responses only (I, green) responses. In grey, no response.

We next asked whether synaptic excitation is followed by inhibition, when few ATN fibers are stimulated to mimic sparse *in vivo* head directional inputs? Rather than a wide-field stimulation, we used a focused blue laser to stimulate small sets of ATN fibers. Layer 3 PCs were recorded in voltage clamp at -30 mV, to separate excitatory and inhibitory synaptic current components, and and low intensity laser scanning photostimulation was randomly applied to different sites across the presubiculum (Fig. 3E-G). Laser stimuli often initiated an inward current followed by an outward current, similar to sequences evoked by whole-field LED stimuli (Fig. 3Ga). This was the case especially for stimulation sites close to the soma of recorded layer 3 PCs. Amplitudes of both inward and outward currents tended to be higher when stimulating in close proximity to the soma, probably due to the activation of a greater number of thalamic fibers that make synaptic contacts. However, laser stimulation could also evoke either only inward (Fig. 3Gb) or only outward currents (Fig. 3Gc). Outward currents were abolished by Gabazine (10 μM, n = 3 out of 3 tested cells) confirming their GABAergic nature, and both inward and outward currents were suppressed by Glutamate receptor antagonists (NBQX 10 μM, APV 100 μM). Overall, for all stimulus sites tested, the probability to evoke sequences of EPSCs and IPSCs was 36 ± 6% (n = 7 neurons from 3 mice), the probability of initiating only an EPSC was 29 ± 3% and that for initiating exclusively an IPSC was 22 ± 3%. For low intensity laser light stimulation of thalamic fibers at different stimulation sites, 13 ± 4% sites did not evoke any synaptic responses in PCs (Fig. 3H). Sparse activation of ATN inputs therefore often, but not always, induces sequential excitatory and inhibitory synaptic events.

### Action potential timing in pyramidal cells and interneurons

We next compared the timing of firing induced in presubicular pyramidal cells (PCs) and parvalbumin (PV) and somatostatin (SST) expressing interneurons by ATN stimulation, to ask ask how the two types of interneuron contribute to disynaptic feed-forward inhibition. (Fig. 4). Action potential latency varied with light intensity: higher intensity stimulation recruited more ATN fibres and induced significantly shorter action potential latencies (6 PV interneurons (green) from 3 mice; 5 PCs (black) from 4 mice, 5 SST interneurons (purple) from 3 mice; p < 0.05, Wilcoxon matched-pairs signed rank test; Fig. 4A). In subsequent experiments, light-intensity was reduced to the minimum value at which action potentials were still initiated in simultaneously recorded PC–PV or PC-SST neuron pairs. In PC–PV pairs, the latency to the first action potential was always shorter for PV interneurons than for pyramidal cells (PC, 3.26 ± 0.26 ms; PV, 1.84 ± 0.44 ms, mean ± sem, n = 7 pairs from 4 mice, p < 0.01, Wilcoxon matched-pairs signed rank test, Fig. 4B,D). In contrast, in PC-SST pairs, pyramidal cells always fired before SST interneurons (latencies: PC, 3.68 ± 0.22 ms; SST, 6.2 ± 0.27 ms, mean ± sem, n = 6 pairs from 4 mice, p < 0.01, Wilcoxon matched-pairs signed rank test, Fig. 4C,E). These data suggest that PV, but not SST interneurons, may mediate thalamic feed-forward inhibition.

**Figure 4:**
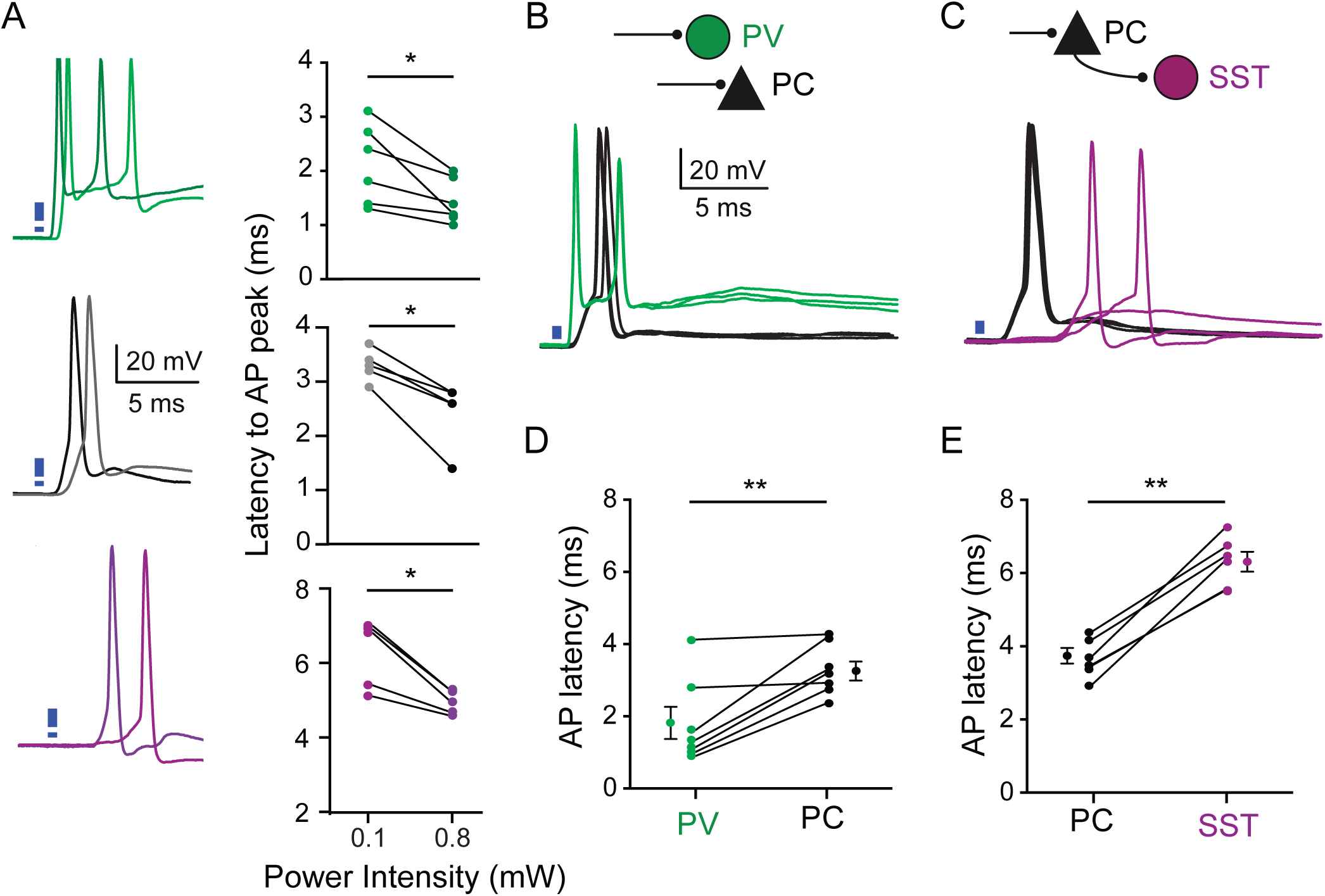
Relative action potential (AP) timing in pairs of pyramidal neurons and interneurons. (A) Left, current-clamp recording of a PV interneuron (green), a pyramidal cell (PC) (black) and SST interneuron (purple), with two different light intensities for stimulation of ATN axon terminals (0.2 and 1.1 mW). Right, summary data. Higher stimulus intensities led to shorter AP latencies in a same neuron. (B) Action potentials initiated by light activation of thalamic fibers in a simultaneously recorded PC–PV pair, or (C) in a PC-SST pair. (D-E) AP latencies from 7 PC–PV pairs, and from 6 PC-SST pairs. Each dot represents 1 neuron. AP latencies in PC cells were longer than in PV interneurons but shorter than in SST interneurons. * p < 0.05, ** p < 0.01, Wilcoxon matched-pairs signed rank test.

### ATN provides direct excitatory inputs to pyramidal neurons and PV interneurons but not to SST interneurons

We next compared responses of presubicular pyramidal cells, fast-spiking PV interneurons and low-threshold spiking SST interneurons to photostimulation of ATN fibers. Pyramidal neuron-interneuron pairs were recorded in presubicular layer 3 in Pvalb-Cre::tdTomato or Sst-Cre::tdTomato transgenic mice (Fig. 5A-F; cf. Nassar et al. 2015). Intrinsic electrophysiological properties of both recorded cells were characterized from hyperpolarizing and depolarizing current pulses (Table 1). Optical stimulation of ATN inputs was adjusted to elicit synaptic excitation in at least one neuron of the recorded pair.

**Figure 5:**
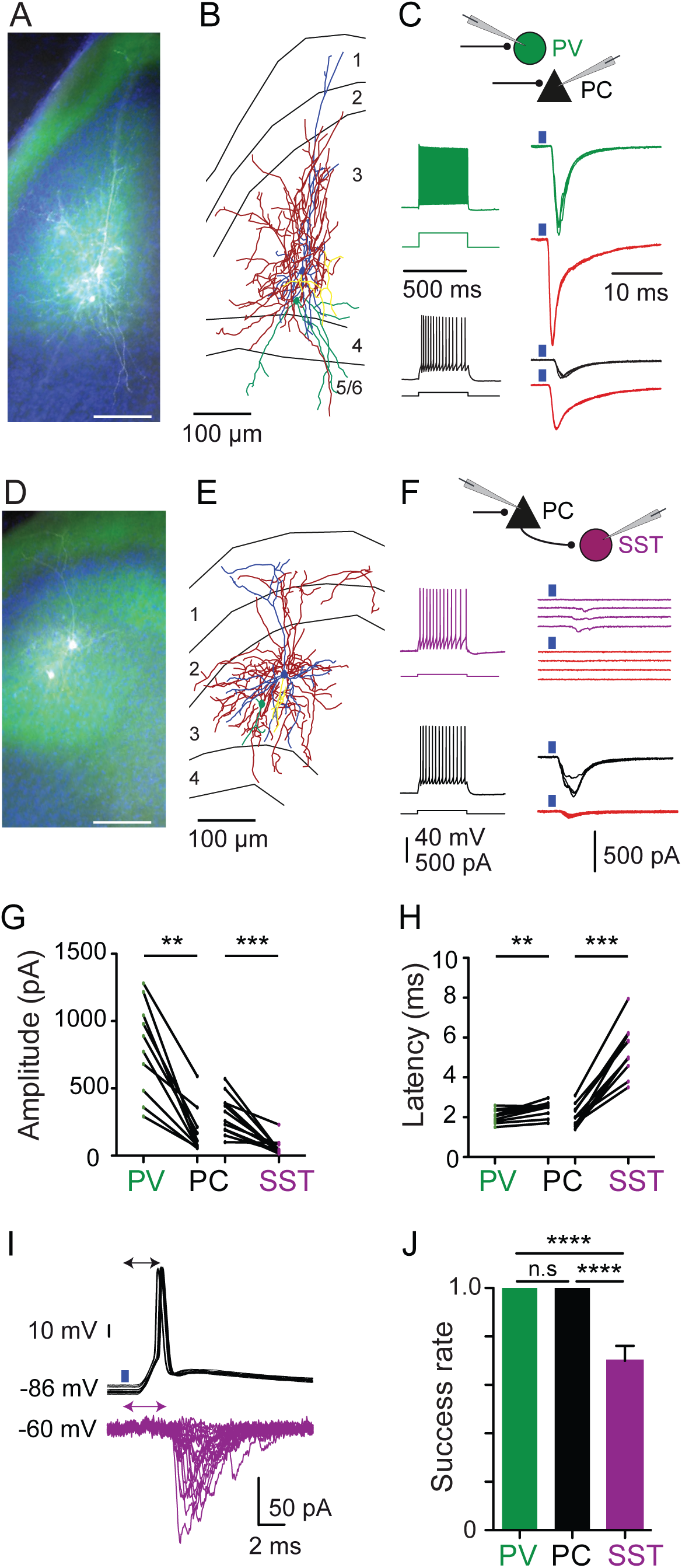
Long-range excitatory inputs from ATN to pairs of PV or SST interneurons and pyramidal neurons in layer 3. (A), (D) Presubicular slice with two biocytin-filled neurons ((A-B), pyramidal neuron and PV interneuron; (D-E), pyramidal neuron and SST interneuron). ATN axons (green) in the superficial layers; scale bars, 100 μm (B), (E) reconstructed cell pairs. Pyramidal cell dendrites in blue, axons in yellow. Interneuron dendrites in green, axons in red. (C), Dual records of PV interneuron (green) and pyramidal neuron (black) or (F), SST interneuron (purple) and pyramidal neuron (black) respectively. Left, firing pattern in response to a two-fold rheobase current injection. Right, light-evoked EPSCs from the illustrated pair of simultaneously recorded pyramidal neuron and interneuron. Red traces, in the prescence of TTX (1 μM) and 4-AP (100 μM) in the bath solution. (G) Amplitudes and (H) latencies from double-recorded principal neuron and interneuron EPSCs. ** p < 0.01, *** p < 0.001, Wilcoxon matched-pairs signed rank test. (I) Light stimulation reliably generated action potentials in a PC. EPSCs in a simultaneously recorded SST interneuron had a 30 % failure rate (30 trials). (J) Bar plots of success rate for synaptic events in different cell types (calculated from 30 trials for each neuron tested; 13 PV interneurons (green), 13 PC (black) and 11 SST interneurons (purple)). **** p < 0.0001, Kruskal–Wallis and Dunn’s multiple comparison post hoc test. Values are given as mean ± SEM. ns: non-significant.

Amplitudes of EPSCs in PV interneurons were significantly larger than those of EPSCs in pyramidal cells (PV, 800 ± 109 pA, n = 10 neurons from 5 animals; PC, 178 ± 55 pA, n = 10 neurons from 5 mice; p < 0.001, Wilcoxon matched-pairs signed rank test, Fig. 5A-C, G). In contrast, the amplitudes of EPSCs in SST interneurons were significantly smaller than those recorded simultaneously in pyramidal cells (PC, 306 ± 41 pA; SST, 58 ± 17 pA; mean ± sem; n = 11 neurons from 10 mice; p < 0.001 Wilcoxon matched-pairs signed rank test; Fig. 5D-G). EPSC onset latencies were similar in PV cells and pyramidal cells (PV, 2.0 ± 0.1 ms; PC, 2.4 ± 0.1ms; mean ± sem; n = 10 neurons from 5 animals; Fig 5H), but significantly longer in SST cells (PC, 2.0 ± 0.2 ms; SST, 5.3 ± 0.4 ms; mean ± sem; n = 11 recordings from 10 animals, p < 0.001, Wilcoxon matched-pairs signed rank test; Fig. 5H). Light-stimulation of ATN axons reliably evoked EPSCs in all pyramidal cells and PV interneurons, but not in SST interneurons. While SST cells responded at least once to repeated trials, failures occurred with a 30% probability across trials (success rate, 0.70 ± 0.06; mean ± sem; n = 11 neurons from 6 animals, Fig. 5I-J, p < 0 .0001, Kruskal–Wallis and Dunn’s multiple comparison *post hoc* test). These results were obtained in conditions where presynaptic action potentials evoked synaptic events. Evidence for direct monosynaptic inputs to pyramidal neurons and PV interneurons was confirmed by showing that synaptic responses persisted during TTX and 4-AP perfusion (1 μM TTX and 100 μM 4-AP; n = 5 pairs from 5 animals; Fig. 5C red traces). However, evoked responses in SST interneurons were abolished after perfusion of TTX and 4-AP (Fig. 5F, red traces), compatible with an indirect recruitment of SST neurons, following thalamic PC activation. Thus, ATN fibers directly excite pyramidal neurons and PV interneurons but not SST interneurons. PV cells therefore appear well-suited to mediate thalamic feed-forward inhibition onto PC cells.

### Presubicular PV interneurons are highly interconnected with pyramidal neurons

We next examined interactions between PV interneurons and layer 3 pyramidal cells in double recordings from synaptically connected cell pairs (n = 40). Figure 6 shows typical morphologies and synaptic interactions of a reciprocally connected PV-PC pair. Action currents initiated in PV cells elicited IPSCs of monosynaptic latencies (0.68 ± 0.03 ms, n = 19, Table 2) in the pyramidal cell (Fig. 6B; n = 20 of 40 tested pairs. Conversely, pyramidal cell action currents initiated unitary excitatory postsynaptic currents (uEPSCs) with monosynaptic latencies (0.74 ± 0.03 ms, n = 17, Table 2; n = 21 of 40 tested pairs). Synaptic interactions between PV interneurons and pyramidal cells were often reciprocal (n = 15 of 40 tested pairs, Fig. 6B), suggesting a high portion of PV cells mediate both feed-forward and feed-back inhibition. IPSCs had rapid kinetics consistent with inhibitory synapses terminating on peri-somatic sites on pyramidal cells. The rise time of unitary IPSCs was 0.52 ± 0.07 ms (n = 19). The rise time of unitary EPSCs from PC-to-PV cells was 0.31 ± 0.02 ms (n = 17). The mean amplitude of EPSCs induced in PV cells was 73 ± 20 pA (n = 17; Fig. 6C, E) and their mean decay time constant was 0.91 ± 0.07 ms (n = 17). The mean amplitude of IPSCs elicited in pyramidal cells was 14 ± 2 pA (n = 19; Fig. 6C, E) and their mean decay time constant was 2.90 ± 0.20 ms (n = 19). The transfer rate, or probability that a postsynaptic event was induced, was 0.79 ± 0.06 for EPSCs (n = 17) and 0.71 ± 0.07 for IPSCs (n = 19). During 50 Hz trains of action currents initated in pyramidal cells or in interneurons (Fig. 6D-F), EPSC amplitudes became moderately depressed (paired pulse ratio, 0.82 ± 0.07, n = 15) as were those of IPSCs (paired pulse ratio, 0.85 ± 0.03, n = 18). We conclude that layer 3 presubicular PV cells mediate a widespread inhibition of local pyramidal cells.

**Figure 6:**
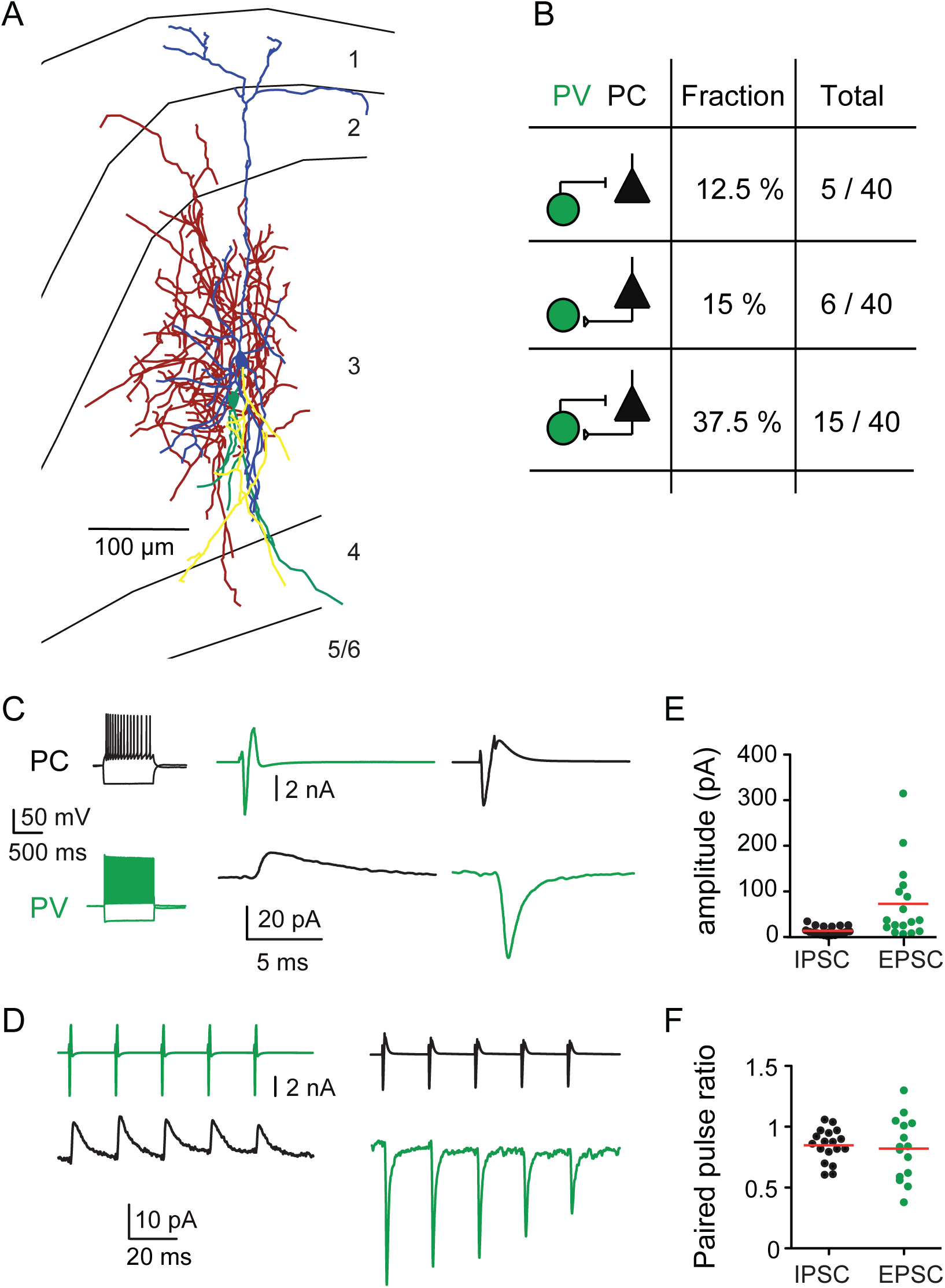
Presubicular layer 3 PV interneurons and pyramidal cells are highly interconnected. (A) Anatomical reconstruction of a pair of reciprocally connected PV interneuron and pyramidal neuron in presubicular layer 3. Pyramidal cell dendrites in blue, axon in yellow; PV dendrites in green, axon in red. Subiculum is to the left and the parasubiculum to the right. (B) Summary of connectivity between PV and PC cells. (C) Paired recording of PV-PC pair shown above. Left, firing patterns in response to a negative and positive (two-fold rheobase) current injection. An evoked action current in the PV cell (green, -70 mV, voltage clamp) initiated a short-latency IPSC in the pyramidal cell (black, -50 mV, voltage clamp). Conversely, an evoked action current in the pyramidal cell elicited an EPSC in the PV cell. Traces are averaged from 10 trials. (D) Unitary IPSCs (black) and EPSCs (green) from the same cell pair induced by 5 action currents at 50 Hz. (E) Absolute amplitudes and (F) paired pulse ratio of IPSCs in PC neurons (black) and EPSCs in PV interneurons (green). Each dot represents averaged recordings from one neuron.

**Table 2.** EPSC and IPSC properties from synaptically connected PV interneuron-pyramidal neuron pairs.

|  | EPSCs |  |  | IPSC |  |  |
| --- | --- | --- | --- | --- | --- | --- |
|  | mean | sem | N | mean | sem | N |
| Rise time (ms) | 0.31 | 0.02 | 17 | 0.52 | 0.07 | 19 |
| Decay time (ms) | 0.91 | 0.07 | 17 | 2.90 | 0.20 | 19 |
| Latency Onset (ms) | 0.74 | 0.03 | 17 | 0.68 | 0.03 | 19 |
| Transfer rate | 0.79 | 0.06 | 17 | 0.71 | 0.07 | 19 |

### ATN fibres induce PV interneuron mediated feedforward inhibition in the presubiculum

These data show that ATN axons excite both presubicular layer 3 PV interneurons and MEc-projecting pyramidal cells (Fig. 2 to 5), and that these two cell types are highly interconnected (Fig. 6). We next asked whether PV cells mediate a thalamic feedforward inhibition. To do so, AAV.hSyn.hChR2-eYFP was injected into the ATN and AAV.Ef1a.DIO.eNpHR3.0-eYFP into the presubiculum of Pvalb-Cre mice. This double optogenetic strategy let us test the effect of inhibiting presubicular PV cells on synaptic responses induced in pyramidal cells by ATN fiber stimulation (Fig. 7A). Expression of the light-gated, chloride-pumping halorhodopsin (eNpHR3.0) hyperpolarized presubicular PV interneurons in response to yellow light (Fig. 7B, n = 5 cells from 3 animals). In cell-attached recordings, PV cells fired in response to blue light stimulation of ATN fibers, and firing was inhibited during concomitant activation of eNpHR3.0 (Fig. 7C). In voltage-clamp recordings from layer 3 pyramidal neurons, ATN stimulation induced a sequence of inward (EPSC) and outward (IPSC) currents at 0 mV. Yellow light silencing of PV interneurons dramatically, and reversibly, reduced the feedforward inhibitory current (n = 8 PCs from 3 mice, p < 0.01, Kruskal–Wallis and Dunn’s multiple comparison *post hoc* test) (Fig. 7D,E). As a control, we noted that the peak amplitude of ATN-driven EPSCs was unchanged during yellow light stimulation (-60 mV; light off/light on/light off, n = 8 cells from 3 animals, Kruskal–Wallis and Dunn’s multiple comparison *post hoc* test, Fig. 7F). These data show PV interneurons mediate a thalamic feedforward inhibition of layer 3 pyramidal cells.

**Figure 7:**
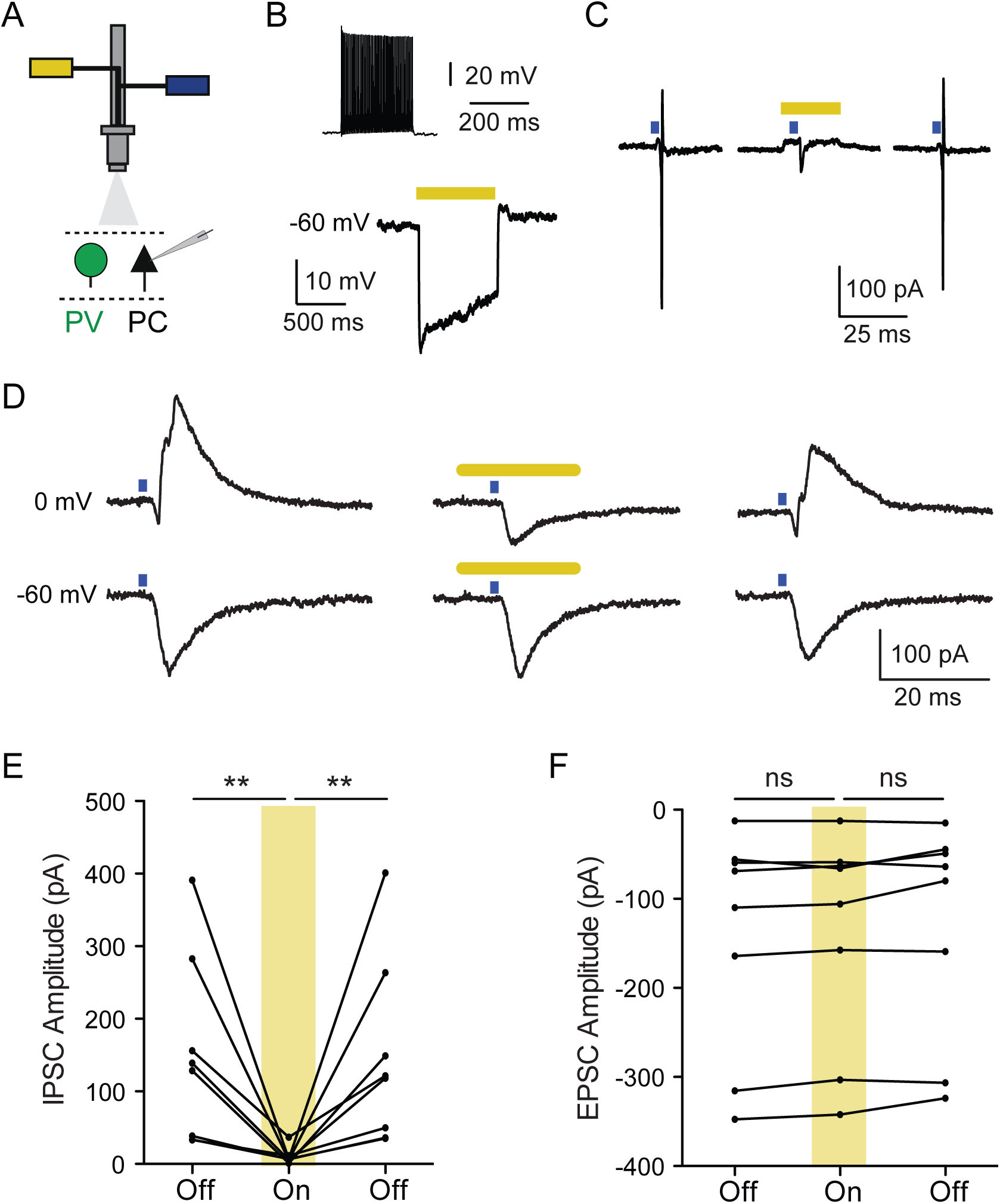
PV interneurons mediate ATN driven feed-forward inhibition in the Presubiculum. (A) Blue (470 nm) and yellow (585 nm) illumination via the same light path was used to activate ChR2 alone, or in combination with eNpHR. (B) A depolarising current step produced high frequency firing in a eNpHR-expressing PV cell. Yellow light activated eNpHR and strongly hyperpolarized this interneuron. (C) Action currents recorded in cell-attached mode from an eNpHR-expressing PV cell in response to photoactivation of ATN inputs with blue light, before, during, and after illumination with yellow light. Yellow light was triggered 5 ms before the onset of the 0.5 ms blue light pulse and remained on for 20 ms. (D) Synaptic currents induced in a layer 3 pyramidal neuron by photostimulation (blue) of ATN fibers in the absence, in the presence, and again in the absence of yellow light. Yellow light stimuli (middle) reversibly inhibited PV interneurons and suppressed evoked IPSCs recorded as outward currents at 0 mV holding potential. (E) IPSCs were suppressed when PV interneurons were silenced, ** p < 0.01, Kruskal–Wallis and Dunn’s multiple comparison post hoc test. (F) EPSCs recorded in PC cells at -60 mV holding potential were not affected by yellow light stimuli. ns: non-significant.

### Effects of repetitive activation of ATN inputs depend on postsynaptic cell identity

We then investigated the dynamics of excitatory postsynaptic current responses to repetitive, low intensity (0.1 - 0.5 mW) stimulation of ATN fibers at 10 or 30 Hz. In PV and PC neurons, the amplitude of light-evoked EPSCs was reduced during trains of ten stimuli at either 10 or 30 Hz (Fig. 8A-F). EPSCs in PV interneurons depressed more rapidly and more strongly (paired pulse ratio (PPR), 10 Hz, 0.63 ± 0.03; 30 Hz, 0.58 ± 0.04, n = 12, mean ± sem; significant depression of EPSC amplitudes after the 3^rd^ stimulus; Anderson-Darling and Hochberg-Benjamini multiple comparison test) than those in pyramidal neurons (PPR, 10 Hz: 0.93 ± 0.04, n = 26; 30 Hz: 0.96 ± 0.05, n = 25, mean ± sem; significant depression after the 6^th^ stimulus; Fig. 8C,F). In contrast, we observed a transient, short-term facilitation of di-synaptic EPSCs in SST interneurons. During stimulus trains at 30 Hz, EPSC amplitude increased significantly between the 1^st^ and the 2^nd^ and 3^rd^ stimulus (PPR, 30 Hz: 5.3 ± 1.19, n = 15, mean ± sem; Fig. 8H,I). Repetitive stimulation at 10 Hz induced very small or no facilitation in SST interneurons (PPR, 10 Hz: 1.56 ± 0.32, n = 14; Fig. 8G,I). These data indicate distinct synapse dynamics for different cell types. Dynamic depression of EPSCs in PV cells should weaken feedforward inhibition of PCs and facilitate firing during maintained head direction signals from the ATN.

**Figure 8:**
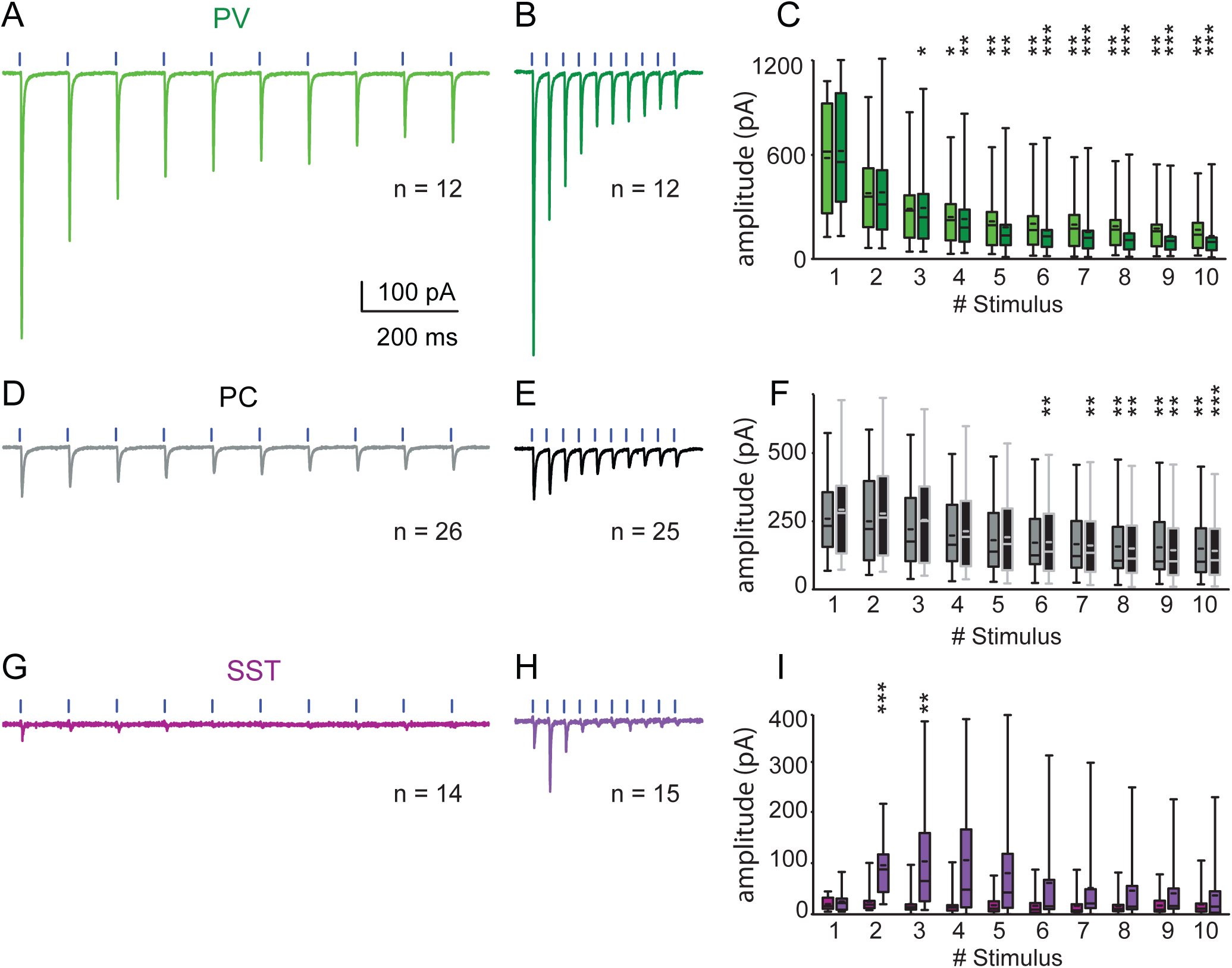
Synaptic dynamics of responses to photostimulation of ATN inputs in layer 3 pyramidal neurons and interneurons. (A), (D), (G) EPSCs evoked by ten successive light stimuli repeated at 10 Hz and (B), (E), (H) at 30 Hz in PV interneurons (light green, dark green), PC cells (grey, black) and SST interneurons (purple, violet). (C), (F), (I) EPSC amplitudes to stimuli at 10 Hz and 30 Hz for the three cell types respectively. Box plots give minimum, maximum, lower and upper quartiles, median and mean values. * p < 0.05, ** p < 0.01, *** p < 0.001, Anderson-Darling test and Hochberg-Benjamini multiple comparison test.

### Dynamics of spiking probability for pyramidal neurons and interneurons

How do differences in synaptic input dynamics affect firing in excitatory and inhibitory presubicular neurons? We examined the evolution of spiking probability during repetitive stimuli. Action potential firing probability was first determined over a range of light intensities (Fig. 7A). In PV interneurons, small increases in light intensity rapidly led to a spiking probability of 1. At higher intensities, single light pulses could initiate more than one AP, thus producing firing probabilities higher than 1, for 12 out of 13 PV cells tested (Fig. 9A). Overall, the probability of action potential discharge was much lower for pyramidal neurons and SST interneurons, for both 10 Hz (PC, 0.5 ± 0.1, n = 22; SST, 0.3 ± 0.1, n = 15; mean ± sem) and 30 Hz stimulations (PC, 0.5 ± 0.1, n = 22; SST, 0.4 ± 0.1, n = 14; mean ± sem) than for PV interneurons (10 Hz, 1.5 ± 0.1, n = 13; 30 Hz, 1.4 ± 0.1, n = 11; mean ± sem). Most PC and SST neurons did not sustain firing until the 10^th^ pulse in a train, even for the higher light intensities.

**Figure 9:**
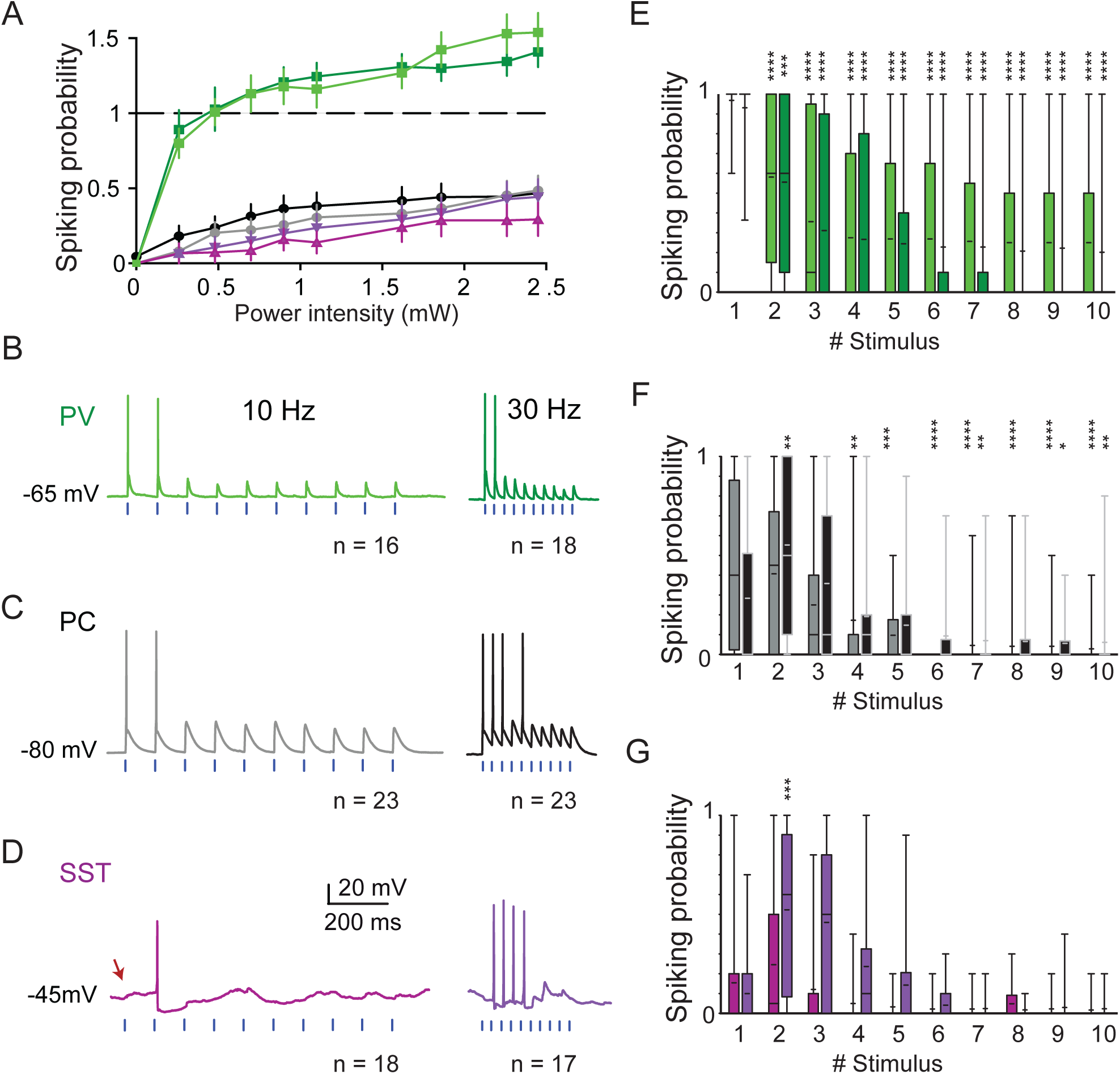
Cell-type specific spiking probabilities following optical stimulation of ChR2 expressing ATN fibers. (A) Variation of spiking probability as a function of light intensity. A spiking probability of 1 corresponds to 10 spikes for a train of 10 stimuli. At higher intensities, a single pulse sometimes induced multiple spikes in PV interneurons (green). PC neurons, black; SST interneurons, purple. Values are given as mean ± SEM. Examples of responses of a PV interneuron (B) pyramidal neuron (C) SST interneuron (D) to 10 Hz and 30 Hz stimulation at resting potential or near AP threshold. Red arrow in (D) indicates the absence of a spike for the 1st pulse. (E), (F), (G) AP spiking probabilities for each cell type for 10 Hz and 30 Hz represented by box plots showing the lower and upper quartiles, the median and the mean. Anderson-Darling test and Hochberg-Benjamini multiple comparison test, to compare the spiking probability of each pulse to the first pulse. * p < 0.05, ** p < 0.01, *** p < 0.001, **** p < 0.0001.

We next compared the dynamics of firing during repetitive ATN fiber stimuli of intermediate light intensities (0.2 to 1 mW). These stimuli evoked at least one action potential but with probability of firing less than 100 % to each pulse. The probability to initiate PV interneuron firing was highest for the 1^st^ pulse of both 10 Hz (n = 16 neurons from 7 animals) and 30 Hz trains (n = 18 neurons from 7 animals, Fig. 9B,E). The spiking probability was then quickly and significantly reduced for the 2^nd^ pulse and during the remainder of the train (p < 0.0001, Anderson-Darling test and Hochberg-Benjamini multiple comparison test, Fig. 9E). The dynamics of firing in PCs differed. At 10 Hz, spiking probability was highest for the 1^st^ pulse, while it was highest for the 2^nd^ pulse at 30 Hz (PC, n = 23 neurons from 10 animals, statistical significance p < 0.01, Fig. 9C, F). After the 4^th^ pulse, and for the remainder of 10 Hz and 30 Hz trains, spiking probability was less than for the first pulse (Fig. 9E, F). Pyramidal neurons and, to a greater extent, SST interneurons did not always fire at the 1^st^ pulse of stimulus trains. In SST cells, we detected a tendency for spiking facilitation for the 2^nd^ pulse for 10 Hz stimulation (SST, n = 18 neurons from 7 animals), and an initial, significant facilitation for 30 Hz stimulation (SST, n = 17 neurons from 6 animals; p < 0.001, Anderson-Darling test and Hochberg-Benjamini multiple comparison test, Fig. 9D, G).

These data reveal a strong initial recruitment of PV interneurons by ATN inputs which depresses during maintained stimuli. Di-synaptically induced firing in SST cells is shaped by facilitating dynamics of the PC-SST synapse and a moderately decreasing probability of PC firing. For stimulations repeated at 30 Hz, the summation of EPSPs and depression of feed-forward inhibition leads to a highest PC spiking probability on the second pulse. We note that thalamic neurons anticipate head direction signals by 25-40 ms (Blair and Sharp 1995; Taube and Muller 1998), while presubicular neurons signal the current direction. Possibly, the fast, transient feed-forward inhibition mediated via PV interneurons helps transform anticipatory direction signals of the thalamus into a real-time head direction signal in presubiculum.

## DISCUSSION

Here we show that glutamatergic synaptic inputs to the presubiculum from the anterior thalamic nuclei (ATN) excite pyramidal cells and PV interneurons, which mediate disynaptic feed-forward inhibition. PV interneurons are recruited at low stimulus intensities by synapses with depressing dynamics. Controlled by inhibitory circuits that are most effective at stimulus onset, pyramidal cell firing increases transiently during repetitive stimulation even though ATN-fiber mediated excitation of principal cells also follows depressing dynamics. In contrast to PV cells, SST interneurons were excited indirectly via pathways that facilitated during repetitive stimulation and so mediate a delayed inhibition during repetitive ATN activity. We showed that activating different sets of ATN fibres in a slice did not always evoke an inhibitory response and that PV interneurons could mediate a lateral inhibition of principal neurons which were not directly excited. These data define excitatory and inhibitory circuits that shape head directional inputs from the thalamus on entry to the presubiculum.

### Optical activation of long-range axon terminals with ChR2

The presubiculum is innervated by thalamus, subiculum, parasubiculum, retrosplenial, visual and entorhinal cortex (Van Groen and Wyss, 1990a,b,c; Van Groen et al., 1992; Jones et al., 2007; Sugar et al., 2011). Here we optically activated thalamic afferents after injecting an AAV-ChR2-eYFP construct in the ATN. Paired recordings of interneurons and pyramidal cells let us define local connectivity and compare the strength and dynamics of ATN inputs.

Synaptic dynamics derived by optical stimulation of ChR2-containing axons should be interpreted with caution (Cruikshank et al., 2010). We took several measures to avoid artefacts. A mutated version of ChR2 with reduced calcium permeability (Lin et al., 2009: Nagel et al., 2003) was used to avoid Ca-dependent effects on transmitter release and synaptic dynamics (Neher and Sakaba, 2008). Synatic depression may be artefactually enhanced according to AAV serotype. Responses determined with AAV9-mediated ChR2 expression are comparable to those induced by electrical stimuli (Jackman et al., 2014). Our data showed both AAV5 and AAV9 serotypes were associated with short-term depression in principal neurons and PV interneurons. Facilitating dynamics of repetitive stimuli in SST interneurons reflect distinct properties of the PC-SST synapse after PC firing. Data obtained by optical stimuli apparently reflect distinct responses of principal neurons, PV and SST interneurons.

### Anterior thalamic afferents excite presubicular layer 3 PCs and interneurons

Layer 3 presubicular head direction cells project to the MEC (Preston-Ferrer et al., 2016; Van Groen and Wyss, 1990c; Honda and Ishizuka, 2008). We showed ATN inputs induced large EPSPs at short latencies in these cells. Intrinsic electrophysiological properties of labeled MEC projecting neurons were similar to those of unlabeled local pyramidal cells (Table 1). Stimulating ATN fibers at 10 Hz induced pyramidal cell firing which diminished during a train. Stimulation at 30 Hz in contrast revealed a high-pass filtering where firing probability increased transiently, before a sustained reduction.

PV interneurons of layer 3 were also directly excited by thalamic fibers. EPSC latencies in PV interneurons and pyramidal cells were short and similar to those of monosynaptic thalamic inputs to barrel cortex (Cruikshank et al., 2010), and other long-range excitatory cortical inputs (Lee et al., 2013; Haley et al., 2016; Keshavarzi et al., 2014). Latencies were somewhat shorter in PV interneurons than in pyramidal cells (Fig. 5) possibly due to distinct subcellular sites of excitatory synapses. ATN afferents may innervate the soma of PV cells while thalamic fibres innervate dendritic spines at some distance from pyramidal cell somata.

In SST interneurons light-evoked synaptic responses were smaller, with longer latencies and transmission failed frequently. SST interneurons were less likely to fire than PV cells, even though their intrinsic excitability and input resistance are higher and resting potential is more depolarized (Table 1). SST cells were excited di-synaptically via layer 3 pyramidal cells, since EPSCs were suppressed by TTX and 4AP (Fig. 5 and Fig. 10; Simonnet et al., 2017). In somatosensory cortex too, circuits for the excitation of PV cells are functionally stronger that those which excite SST interneurons (Cruikshank et al., 2010; Lee et al., 2013).

**Figure 10:**
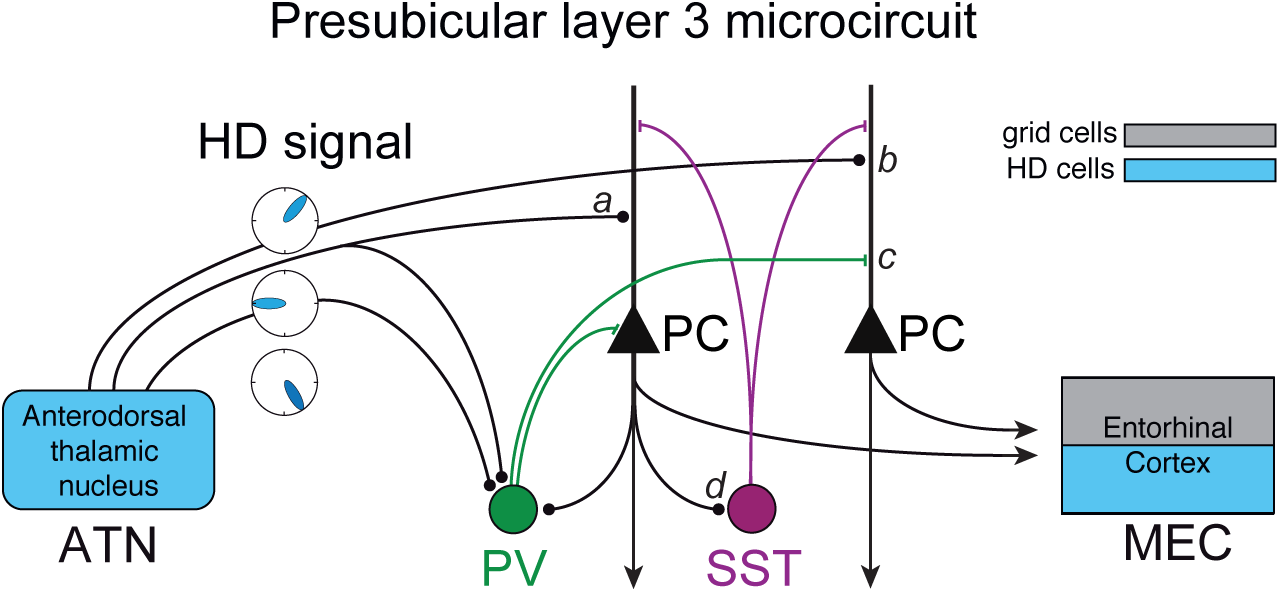
Schematic illustration of connectivity of anterior thalamic (ATN) fibers targeting presubicular layer 3 neurons. Presubicular pyramidal cells (PC) projecting to medial entorhinal cortex (MEC) receive direct excitation from thalamic fibers. Parvalbumin expressing (PV) interneurons also receive direct excitation from thalamic fibers, and they provide transient feed-forward inhibition onto PCs. The activation of single thalamic fibers may either lead to (a) PC excitation followed by feed-forward inhibition, or (b) only excitation, or (c) only rapid disynaptic inhibition of PCs (cf. Fig. 3). SST interneurons (d) are recruited in a feed-back manner following action potential discharge of local PCs (Fig. 4) at high frequency (Fig. 9). SST-PC connectivity is high (cf. Simonnet et al., 2017), and provides activity-dependent feed-back or lateral inhibition.

### Thalamic excitation may be fine-tuned by PV mediated inhibition

Optical stimulation of ATN fibers induced EPSC-IPSC sequences in layer 3 pyramidal cells. IPSC onset latency was short (Fig. 3), as for other thalamocortical pathways (Pouille and Scanziani, 2001; Gabernet et al., 2005; Cruikshank et al., 2010; Stokes and Isaacson, 2010). Our data suggests feed-forward inhibition was largely mediated by presubicular PV interneurons. PV cells fire rapidly when thalamic fibers are stimulated (Fig. 4,5) and contact local pyramidal cells with high probability (Fig. 6). Presubicular PV cells innervated 20 out of 40 pyramidal cells tested, similar to the connectivity of fast-spiking PV cells in somatosensory or visual cortex (Holmgren et al., 2003; Yoshimura and Callaway, 2005). Reciprocal connectivity in the feedback inhibitory circuit was high (Holmgren et al., 2003; Yoshimura and Callaway, 2005; Yoshimura et al., 2005; Packer and Yuste, 2011). Further, optical activation of halorhodopsin let us demonstrate the essential role of PV interneurons in disynaptic feed-forward inhibition (Fig. 7). Silencing presubicular PV interneurons, largely suppressed disynaptic inhibition in pyramidal neurons. PV cells thus mediate a fast, reliable feed-forward control of ATN signals (Fig. 10).

Feed-forward inhibition seems likely to increase with the number of active afferents (Isaacson & Scanziani, 2011). This point was demonstrated by comparing whole-field optical and focal laser stimulation of ChR2 expressing ATN fibers. Whole field stimuli induced balanced EPSC-IPSC sequences in pyramidal cells, but in responses to a more sparse fiber activation induced by focal stimuli (Fig. 3), EPSCs were not succeeded by IPSCs for 20-30% of presubicular sites tested. Possibly inhibition mediated by distal dendritic synapses may have been underestimated due to insufficient space clamp (Williams and Mitchell 2008; Kubota et al., 2016). Indeed it is difficult to compare dendritic excitation and inhibition without making assumptions on the site and strength of contacts (Chadderton et al., 2014). Nevertheless, our data reveals a variable somatic balance of excitation and inhibition which presumably affects synaptic integration and action potential initiation (Markram et al., 2004). We also noted in all cells that disynaptic IPSCs were induced with no preceding EPSC from some stimulation sites. PV interneurons with widespread axonal arbors may thus mediate both feed-forward and lateral disynaptic presubicular inhibition (Fig. 10).

### Functional implications for presubicular HD signal processing

Presubicular layer 3 is a crucial node in head direction signal transmission from the ATN to medial entorhinal cortex. Inputs to the presubiculum probably possess some direction-tuning (Peyrache et al., 2015; Taube, 2007). The feed-forward excitatory-inhibitory microcircuits of layer 3 should sharpen head direction signals relayed to the entorhinal grid cell system (Tukker et al., 2015; Winter et al., 2015; Preston-Ferrer et al., 2016). As for inhibitory circuits of auditory cortex (Wu et al., 2008; Chadderton et al., 2014), our data suggest that head direction cells coding for non-preferred directions may receive inhibitory but not excitatory inputs. As in barrel cortex, direction-dependent shifts in timing of excitation and inhibition may result in a ‘window of opportunity’ for excitatory inputs to induce firing (Wilent and Contreras, 2005). Operations performed in presubicular layer 3 circuits seem well-adapted for spatial fine-tuning of head direction signals sent to the MEC.

Fast-spiking interneuron firing is modulated by angular velocity during head rotation (Preston-Ferrer et al., 2016). Our data shows thalamic excitation of PV cells depresses during repetitive thalamic activity. During a fast head turn however active thalamic afferent fibers change rapidly, so that thalamic excitation, and firing, of PV cells should be maintained. The depressing dynamics of PV cell excitation may thus permit ‘re-construction’ of angular velocity signals by decoding thalamic head directional inputs.

PV interneuron mediated inhibition should be strong during fast head turns. In contrast, when direction is maintained, accompanied by activity in the same set of thalamic fibers, synaptic excitation of PV cells will be depressed. Feed-forward inhibition should weaken as PV cells fire less and the excitatory-inhibitory balance may become shifted to excitation (Whitmire and Stanley, 2016) if synaptic depression is greater for inhibition than for excitation, as in barrel cortex (Gabernet et al., 2005).

As thalamic head direction signals are initiated, feed-forward inhibition in the presubiculum may override excitatory signals. For a maintained head direction, depressing dynamics of the ATN-PV synapse will favor delayed pyramidal cell recruitment, by enhancing thalamic excitation for later stimuli. Thus, PV interneuron circuit properties permit both fine-tuning of rapidly changing ATN inputs and a distinct treatment of persistent inputs.

During immobility, pyramidal cells excited by ATN inputs will in turn effectively excite SST interneurons due to the facilitating dynamics of the excitatory synapses (Ma et al., 2010). We showed previously that recurrent connectivity is high between SST expressing Martinotti type interneurons and layer 3 pyramidal cells. Precisely timed inhibitory feedback from SST cells occurring during pyramidal cell spike repolarization sustains PC firing (Simonnet et al., 2017). SST cell activity during maintained head direction may then enhance pyramidal cell firing. Presubicular circuits could then switch between two regimes according to the angular velocity of head movements. During immobility, SST-PC interactions support maintained HD firing by attractor-like mechanisms. During rapid head turns, in contrast, PV mediated inhibition acts to tune the HD signal transmitted to MEC.

## ACKNOWLEDGEMENTS

This work was supported by the French Ministry for Education and Research (M.N) and the Centre National des Etudes Spatiales (M.N. & M.B), the Région Ile-de-France (J.S.) and by ANR Grant JCJC R10206DD (D.F.). The research leading to these results also benefitted from the program “Investissements d’avenir” ANR-10-IAIHU-06 and we acknowledge financial support from the ERC (322721, RM). We thank thank Nathalie Sol-Foulon for help with confocal imaging, and Karl Deisseroth for making available pAAV-hSyn-hChR2(H134R)-eYFP and pAAV-Ef1a-DIO eNpHR 3.0-EYFP.

